# Cell-Specific Modulation of the Aryl Hydrocarbon Receptor by Kynurenine in Pulmonary Fibrosis Requires Microenvironmental Crosstalk

**DOI:** 10.64898/2026.07.21.738475

**Authors:** Hannah Carter, Benjamin C. Anderson, Rita de Medina Costa, Joshua Franzen, Joshua Krukonis, Kayla C. Jenkins, Rachel L. Zemens, Bethany B. Moore, Stephen J. Gurczysnki

## Abstract

**Background:** Idiopathic pulmonary fibrosis (IPF) is a progressive, chronic lung disease with limited therapeutic options. Tryptophan metabolism is significantly dysregulated during lung fibrogenesis, with the metabolite kynurenine (kyn) accumulating in lung tissue and driving pathology via the aryl hydrocarbon receptor (AHR). This study evaluates the cell-specific contributions of kyn-mediated AHR signaling across different pulmonary cell types to clarify its role in disease progression.

**Methods:** Using a murine model of bleomycin-induced pulmonary fibrosis, lung tryptophan metabolites were profiled via liquid chromatography-mass spectrometry. The functional and transcriptomic impacts of kyn administration and AHR modulation were subsequently characterized across three distinct cellular compartments: CD103+ dendritic cells (DCs), fibroblasts, and alveolar epithelial cells (AECs).

**Results:** Kyn levels were elevated in fibrotic lungs, and exogenous kyn selectively exacerbated collagen deposition during the fibrogenic phase rather than altering acute injury. *In vitro* monocultures of primary lung fibroblasts and AECs revealed negligible functional responses to kyn or AHR inhibition regarding myofibroblast differentiation, migration, or epithelial barrier disruption. Intriguingly, primary tissue-resident CD103+ DCs exhibited a hyperinflammatory, non-canonical AHR signaling profile *in vivo*. While *ex vivo* monoculture rapidly reverted these DCs to an anti-inflammatory, canonical AHR state, directly co-culturing DCs with fibrotic primary lung fibroblasts successfully restored the pathogenic, non-canonical signaling phenotype characterized by augmented IL-6 production and suppressed canonical targets.

**Conclusions:** Pathogenic AHR signaling in pulmonary fibrosis is highly cell-context dependent and driven by complex cell-cell interactions. Reductionist monocultures fail to replicate tissue-level dendritic cell phenotypes, highlighting the necessity of co-culture models and providing a cautionary note for the systemic clinical use of AHR-targeted therapeutics.

## Introduction

Idiopathic pulmonary fibrosis (IPF) is a progressive, chronic lung disease characterized by scarring of the lung tissue, which affects roughly 3 million people world-wide.(1) IPF has no known cause, but is considered a disease of dysregulated wound healing, characterized by chronic inflammation, aberrant epithelial responses, and overactivated fibroblasts.(2) The average prognosis from the time of diagnosis is only 2-5 years.(3) Although steroids and anti-inflammatory therapies have been ineffective clinically, numerous lines of evidence in both human disease and animal models point to important regulatory and pathogenic roles for immune cell crosstalk with structural cells in lung fibrosis.(4) Thus, a deeper understanding of how different cell types interact to produce pulmonary fibrosis is critical to understanding this disease.

The aryl hydrocarbon receptor (AHR) is a ligand activated transcription factor which has historically been understood to regulate metabolism of environmental pollutants.(5–7) In the last 20 years, however, it has been appreciated that AHR also impacts immune responses, cell differentiation, and the function of many cellular pathways.(8–16) What effect AHR may have on a system depends on the cell type, the AHR ligand involved, and crosstalk with other proteins.(15, 17–21) There is mounting evidence that AHR can participate not only in sensing environmental ligands through the canonical AHR pathway, but also coordinates transcription of inflammatory genes through dimerization with non-canonical transcription factors like NF- κB.(22–24) The Pleotropic nature of AHR signaling has produced seemingly contradictory, nuanced, results regarding its ultimate role as an pro- or anti-inflammatory mediator. Furthermore, AHR is most often studied using strong, exogenous ligands like 2,3,7,8- Tetrachlorodibenzo-p-dioxin (TCDD) which may not be relevant in many disease models. Naturally occurring AHR ligands, like the kynurenine (kyn) family of tryptophan metabolites, have been relatively understudied in their role in participating in AHR driven pathological processes.

There are elevated levels of kyn in the plasma of IPF patients and in the lung tissue of fibrotic mice that have received bleomycin.(25–27) In the context of pulmonary fibrosis, we previously found that treating mice with kyn during fibrogenesis in the blm mouse model led to worse fibrosis, whereas treatment with the AHR antagonist CH223191 was protective.(26) Our results showed that dendritic cells accumulate post-blm, express large amounts of AHR and are necessary for robust fibrogenesis, however, DCs do not undergo canonical AHR signaling in- vivo and instead engage in pro-inflammatory non-canonical AHR signaling.(26) Other studies have also highlighted the importance of non-canonical AHR signaling specifically in DC maturation and activation.(28–30) What factors might drive the dendritic cells toward non-canonical signaling in the lung are not known. Furthermore, the contribution of AHR signaling in other cell types relevant for fibrogenesis such as fibroblasts and epithelial cells remains relatively under-studied. Here, we take a reductionist approach to understand how kyn might impact these cell types. Our results show that complex cell-cell interactions take place in the lung that drive AHR into a pro-inflammatory role. And importantly, *ex vivo* culture conditions cannot easily recapitulate the DC phenotype noted *in vivo*. Thus, caution must be exercised when extrapolating AHR function from ex vivo experiments.

## Material and Methods

### Sex as a biological variable

Our study exclusively examined male mice. IPF is predominantly a disease affecting male patients. In line with this, female mice develop much-less-severe fibrosis in response to bleomycin(31–33).

### Mouse lines and blm

WT C57Bl/6J mice were ordered from Jackson labs and used between 6-8 weeks of age. Bleomycin was purchased as Blenoxane from the pharmacy at the University of Michigan hospital, 1U/mL stocks were made using sterile saline and mice were administered 0.75-1U / kg via the oropharyngeal route. Kynurenine, hydroxykynurenine, indole-propionic acid, and CH223191 were all dissolved in DMSO and administered to mice via oral gavage in a vehicle solution of phosphate-buffered saline with 2% Tween-20 and 0.5% carboxy-methyl cellulose daily on days 10-21 and harvested on day 21 unless otherwise noted. Kynurenine (Thermofisher) was used at 40mg/kg, CH223191 (Tocris) was used at 5mg/kg. Hydroxykynurenine was used at 40mg/kg, and IPA was used at 40mg/kg for *in-vivo* blm experiments. Bronchoalveolar lavage fluid was collected from either saline- or blm- treated mice 3 days after blm by washing 1mL of sterile saline intratracheally through the lungs. Total protein was quantified using BCA assay (Pierce) and IgM through ELISA (Invitrogen) per manufacturer’s instructions. All animal experiments were approved by the U-M Institutional Committee on the Use and Care of Animals under protocol PRO00011303.

### Characterization of lung tryptophan metabolites via mass spec

Mouse lungs were harvested and one lobe flash frozen immediately. Lung samples were weighed and 50-100 mg of lungs transferred to a microtube with 1.0 mm diameter zirconia/silica beads, and 1ml of extraction solvent, 1:1:1:1 methanol:acetonitrile:water:acetone containing isotope labeled internal standards (Tryptoophan-D5, Kynurenic Acid-D5, L-Kynurenine-D4, 5- Hydroxytryptophan-D3, 5-Hydroxyindoleacetic Acid-D6, 3-Hydroxyanthranilic Acid-D3) was added in a ratio of 33.33 μL/mg. Tissue disruption was performed in a FastPrep-24 5G (MP Biomedicals LLC) using the recommended program for mouse lung at room temp (speed: 6 m/s, time: 40 sec in 1 cycle and snap-frozen on dry ice and stored at -80°C. Further sample processing was performed by the Metabolomics core at the University of Michigan, in short, samples were thawed and incubated on ice for 10 minutes and vortexed to remix and centrifuged at 14,000 RPM for 10 min at 4°C. 200 μL of extract was transferred to an autovial and dried under a continuous stream of nitrogen at room temperature for 2-h. Dried samples were reconstituted in 100 μL of 80:20 water/methanol. A pooled sample for quality control was created by combining 20 μL of extract from each sample and processed as the other samples. This sample was run periodically throughout the mass spectrometry analysis. A calibration curve containing 7 standards was prepared alongside the samples. The curve contained the following metabolites and concentration ranges: 3-Hydroxyanthranilic Acid, 3-Indolepropionic Acid, 2-Aminobenzoic Acid (Anthranilic acid), Kynurenic acid, L-Kynurenine, Xanthurenic Acid, Indole-3-lactic acid, Picolinic Acid, Indole-3-acetic acid, Serotonin, all from 0-2 μM and 5- Hydroxytryptophan, 5-Hydroxyindoleacetic acid, L-Tryptophan all from 0.5-40 μM. LC-MS analysis was performed on an Agilent system consisting of an Agilent 1290 Infinity II / 6545 qTOF MS system with the JetStream Ionization (ESI) source (Agilent Technologies, Inc) using an Acquity HSS T3, 2.1 mm x 100 mm, 1.7 µm particle size column (Waters Corporation). Each sample was analyzed twice, once in positive and once in negative ion mode. Mobile phase A was 100% water, B was 100% methanol and C was 2.5% formic acid. The gradient for both positive and negative ion modes was as follows: .0% B, 4% C (0 min), 99% B, 1C% (10 min), 99%B, 1%C (17 min), 0%B, 1%C (17.1min), The column was then reconditioned for 3 min at starting conditions before moving to the next injection. The flow rate was 0.45 mL/min and the column temperature was 55°C. The injection volume for positive and negative mode was 10 µL. Source parameters were: drying gas temperature 275°C, drying gas flow rate 12 L/min, nebulizer pressure 45 psig, sheath gas temp 325°C and flow 12 l/min, and capillary voltage 4000V, with internal reference mass correction. *Data Analysis:* Data were processed using MassHunter Quantitative analysis version B.10.00. Metabolites were normalized to the nearest isotope labeled internal standard and quantitated using 2 replicated injections of 7 standards to create a linear calibration curve with accuracy better than 90% for each standard. Several metabolites could be detected in both positive and negative mode, with quantitation typically agreeing within a 10% coefficient of variation (CV). Final data were reported in the mode for which each metabolite had the greatest signal (area) and the lowest CV in the pooled sample. Quantitative data was normalized to tissue mass and reported as fold change of blm-treated samples relative to saline treated samples.

### Quantitation of lung Hydroxyproline

Lungs from BLM or WT mice were resected at the indicated timepoints and incubated overnight at 110°C in 6N hydrochloric acid in well-sealed glass tubes. The following day, aliquots of acid-hydrolyzed lung homogenate were incubated with a chloramine-T solution and then reacted with 4- (Dimethylamino)benzaldehyde. Hydroxyproline was detected via absorbance at 560 nM on a synergy H1 spectrophotometer (Biotek). This method was described previously.(34)

### Preparation of tissue for histology

Formalin-fixed paraffin-embedded lung sections were prepared as follows. Mice were euthanized by CO_2_ asphyxiation, after which 3–5 ml of phosphate-buffered saline was perfused through the right ventricle of the heart. Lungs were infused with 1 ml 10% formalin solution through the trachea and fixed for 24 h, after which lungs were dehydrated in 70% ethanol, embedded in paraffin and sectioned to a 10 μM thickness. Lung sections were subsequently stained with Masson’s Trichrome stain to highlight collagen fibers.(35)

### IHC staining

Paraffin-embedded slides were gradually rehydrated, then antigen was retrieved by pressure cooking slides for 20 minutes in 10mM sodium citrate buffer with 0.05% Tween-20, pH 6.0. Slides were then washed in tris-base saline with 0.1% Tween-20 (TBST) and blocked in 5% donkey serum for one hour in a humidified chamber. Primary antibodies used were: GFP (Abcam ab13970 used at 0.1mg/mL), CD103 (GeneTex GTX53154 used at 2ug/mL), and CD11c (Sysy Antibodies HS-375 003 used at 1:100 dilution). Slides were washed twice in TBST for five minutes then incubated in secondaries (all from JacksonImmuno Laboratories, 703-096- 155, 712-175-150, and 715-165-150, all at 1:250 dilution) for 1h before being washed again. Slides were incubated with DAPI (1:10,000) for 10 minutes, washed, and mounted in Prolong Gold (ThermoFisher).

### Isolation of RNA for qRT-PCR

RNA quantity and purity was assessed on a nanodrop spectrophotometer (Thermo). Transcript expression was analyzed via reverse transcriptase quantitative real-time PCR using Lunaprobe reagent (New England Biolabs) on a Quantstudio3 thermocycler (ABI). Expression analysis was conducted using a ΔΔct calculation. Transcripts were normalized to expression level of a housekeeping gene, RPL38, using a pre-validated primer probe set (Integrated DNA technologies, Hs.PT.58.40595235.gs). Other primer and probe sequences are detailed in **Table 1**.

### RNAseq data analysis pipeline

RNAseq was conducted as previously described.(8) In short, RNA was processed and sequenced by Genewiz (Azenta) using their standard RNAseq analysis package. Raw transcript reads were normalized and differential gene expression (DEG) analysis was conducted using a combination of R and Python with the DESEQ2 package. Lists of DEGs were filtered against a database of protein coding open reading frames using the Biomart package in Python with a fold change cutoff of > 2.0 and p < 0.05. Significant DEGs were processed through the Enrichr package in Python using the KEGG 2026 database to generate lists of overrepresented functional categories. Over representation analysis was conducted in Python to identify core driver genes that appeared in multiple Enrichr categories with a Combined Score of > 25 (genes found in > 2 or more categories were identified from the down-regulated Enrichr dataset while genes present in > 3 or more categories were identified from the up-regulated Enrichr dataset). Core driver gene lists were then interrogated using the STRING package (stringency 7) with MCL clustering (granularity of 2.5) in Cytoscape 3.10.4 to identify gene interaction networks. Lists of all significant DEGs as well as the complete functional Enrichr dataset are provided in supplemental 1.

### Generation of inducible CD103+ DCs

Bone marrow cells were aseptically isolated from the femurs of indicated mouse lines and incubated for 16-days in RPMI media containing 10% heat inactivated FBS, 1% Pen/Strep cocktail, 50µM β-mercaptoethanol, 3 ng/mL GMCSF (R&D systems) and 200 ng/mL Flt3-L (R&D systems) before any additional treatment. iCD103 cells were characterized to be CD11c+, MHCII+, and CD103+ by flow cytometry.

### Cell cultures and reagents

MLE-12 cells were grown in DMEM:F12 media with 10% fetal bovine serum and 1% Pen/Strep cocktail. For certain experiments MLE-12 cells were grown in DMEM:F12 supplemented with 2% FBS and HITES cocktail (10 nM hydrocortisone, 0.005 mg / mL insulin, 0.01 mg / mL Transferrin, 10 nM β-estradiol, 30 nM sodium selenite) as previously described.(36) CCL210 cells were grown in Minimal essential media with 10% fetal bovine serum and 1% Pen/Strep cocktail. Cells were used between 10 and 40 passages. To cells at ∼80% confluence in cell culture, kyn was given at 200uM and CH223 was given at 20μM. For TGF-β, recombinant mouse TGF-β (R&D Systems) was used for MLE-12 cells and iCD103s, and porcine TGF-β (R&D Systems) was used for CCL210 cells at the doses noted.

### Generation of single cell suspensions for murine DC isolation and flow cytometry

Mouse lungs were first perfused via injection of 3-5mL of saline through the right ventricle of the heart. Lungs were resected and minced with scissors before incubation in a collagenase buffer containing 1 mg/mL Collagenase-A (Roche), 2000U DNaseI (Sigma), in DMEM media supplemented with 10% heat inactivated FBS and 1% Pen/Strep cocktail for 45 minutes at 37°C in a shaking incubator. Cell suspensions were disrupted by drawing through a 10 mL syringe before filtration through a 100 µm pore sized cell strainer. Resulting single cell suspensions were assayed for cell viability via trypan blue exclusion.

CD103+ lung DCs were isolated via a 2-step magnetic bead isolation procedure. Single cell collagenase suspensions were first processed via the MojoSort mouse pan dendritic cell isolation kit (BioLegend) following the manufacturer’s protocol. Resulting crude DC preparations were next processed using a FITC labeled anti-CD103 primary antibody in conjunction with anti-FITC magnetic beads (Stemcell Technologies) following the manufacturer’s protocol.

Staining for flow cytometry was carried out via incubation with appropriately diluted fluorophore conjugated antibodies. Antibodies used for flow cytometry were as follows: Myeloid flow cytometry staining panel, BV650-CD11b (clone M1/70), BV510-CD45 (clone 30-F11), BV421-I- Ab (MHCII clone AF6-120.1), APC-Cy7-SiglecF (clone E50-2440), APC-Ly6G (clone 1A8) purchased form BD Horizon. BV605-CD64 (clone X54-5/7.1), PerCP-Cy5.5-CD24 (clone M1/69), PE-Cy7-Ly6C (clone HK1.4), PE-eFluor610-CD11c (clone N418), PE-AHR (clone 4MEJJ) purchased from eBioscience. Note: Some markers were stained for but were not graphed. Flow analysis was carried out in FlowJo v. 10.5.

### Western blot

4×10^5^ CCL210 cells were plated per well of a 6-well plate in 2mLs of complete MEM. After 24 hours, cells were changed to serum free media overnight, then treated with TGF- β if appropriate. After 24 hours, cells were treated with 200µM Kyn or 20µM CH223191 for two days before being washed once with PBS and collected in RIPA buffer with protease inhibitor (Holt). Lysates were normalized according to total protein, reduced in Laemmli buffer with 8% β- mercaptoethanol, boiled for 5 minutes, then separated in a 4-20% PAGE gel before being transferred to a PVDF membrane. Cells were blotted against α-Smooth Muscle Actin antibody (eBioscience) and GAPDH antibody (Cell Signaling Technologies).

### Fibroblast isolation and scratch wound assay

Fibroblasts were isolated as previously described.(37) In short, groups of mice were administered blm or saline as control. Twenty-one days after blm treatment, lungs were harvested and minced with scissors. Minced lungs were incubated in standard incubator conditions and DMEM with 10% FBS and 1% Pen/Strep cocktail with gentle media changes after 4 days. At 7 days cells were trypsinized and split 1:4 and allowed to grow for an additional 7 days with media changed at day 11. Cells were trypsinized and plated on day 14 at 20,000 cells/well of a 96-well Sartorius ImageLock plate. After adhering for 24 hours, cells were scratched using a Sartortius WoundMaker, washed twice with cold PBS, and left in complete DMEM with DMSO (veh), kyn or CH223191. Cells were imaged using an IncuCyte system over 3 days and analyzed using the Sartorius IncuCyte analysis software.

### Alveolar Epithelial Cell (AEC) isolation and Transepithelial Electrical Resistance (TEER) analysis

Lungs were harvested from healthy mice then digested with dispase (Corning) and DNase (Sigma). After filtering through subsequent 100µm and 40µm strainers, the resulting single cell suspension is negatively selected against erythrocytes, myeloid cells, lymphocytes, and endothelial cells using antibodies against TER119, CD32, CD45, and CD104 (all Biolegend) and straptavadin-bound magnetic beads (Promega). The resulting cell suspension is then positively selected for epithelial cells with a PE-EpCam antibody (Biolegend) using the PE- Releasable Positive Selection kit from StemCell. 2.5×10^5^ cells were plated onto fibronectin-coated, 24-well sized, 0.4µm pore transwells, allowed to adhere for three days in HITES media with 10% FBS. Basal media was changed and apical media was removed to establish air liquid interface for an additional 4 days, with basal media changed at day 2. Fresh media was then added to the apical and basal sides for 24 hours before beginning 7hour TEER challenge with 100ng/mL TNFα and 100ng/mL IFNγ in both conditions, and 200uM kyn in one condition. TEER was measured via World Precision Instruments EVOM^3^ system.

## Results

### Kyn is increased and uniquely drives fibrosis in blm-induced pulmonary fibrosis

We showed previously that giving kyn to mice via oral gavage worsens blm-induced fibrosis(26). Others have evaluated how tryptophan and kyn are altered in the serum and the lung tissue of both patients with IPF and mice with blm-induced fibrosis(25, 38), however the rest of the tryptophan metabolism pathway has been largely overlooked. Tryptophan can be metabolized through the kyn pathway (KP), serotonin pathway, or through bacterial metabolism via various indole metabolites **(Figure 1A and ref** (39)). We measured tryptophan metabolites directly in the lung tissue of blm- or saline- treated mice after 21 days, at peak fibrosis(40) and found that there was a significant increase in the lung tissue of tryptophan, as well as kyn and its immediate metabolite, hydroxykynurenine (HK), and the bacterially-derived metabolite 3- indolepropionic acid (IPA) **(Figure 1B)**. As we have previously found(26), treatment with additional kyn (40mg/kg delivered d10-21 post blm) drives worse fibrosis and blockade of AHR signaling ameliorates fibrosis **(Figure 1C and 1D).** However, treatment with HK (40 mg / kg) or IPA (80 mg / kg) did not drive any change in fibrosis **(Figure 1E and 1F)**. This leads us to conclude that altering tryptophan metabolism and kyn levels affect fibrosis, but not by changing serotonin levels, altering production of other KP metabolites, or altering bacterial tryptophan metabolism.

**Figure 1.**
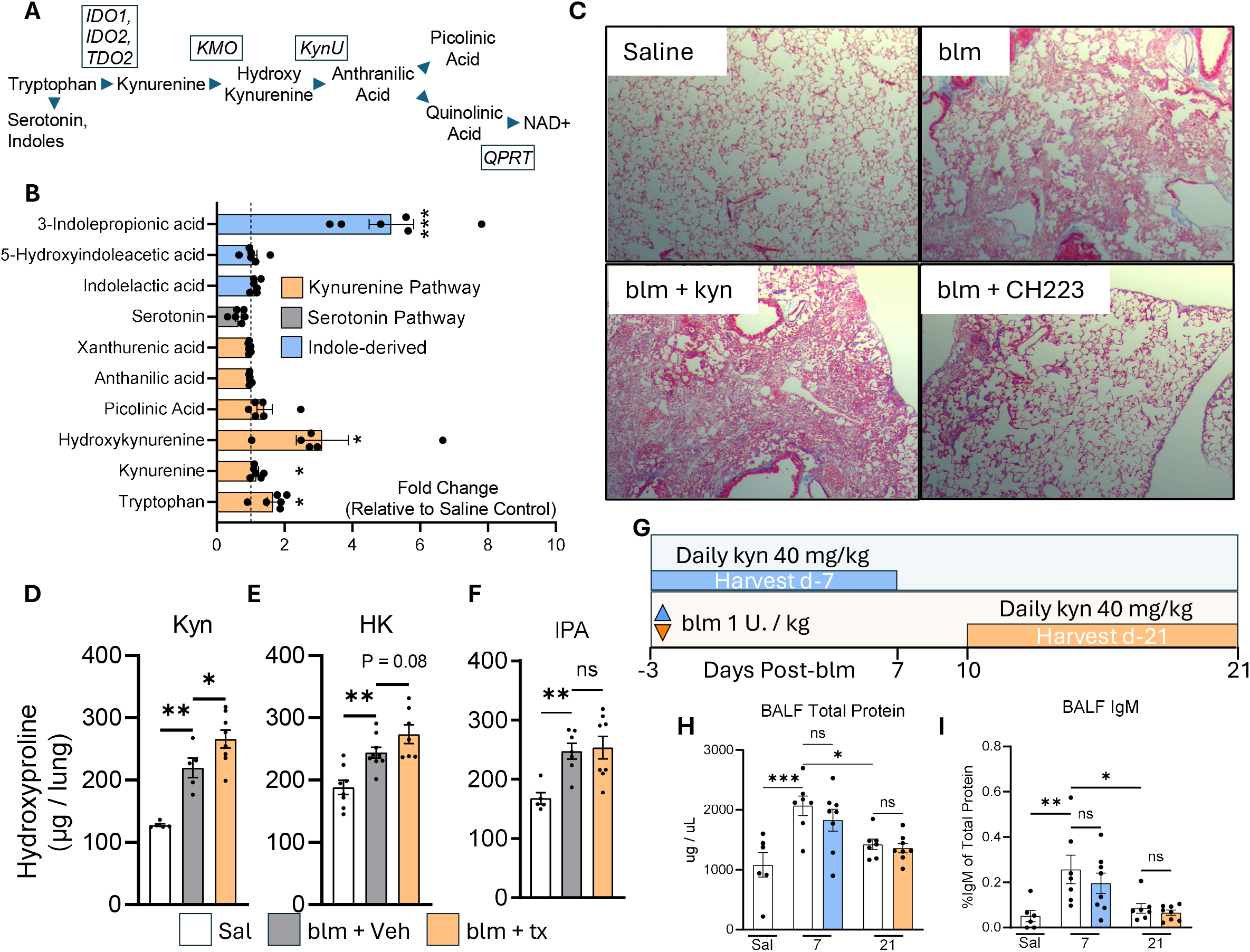
Tryptophan metabolism is altered in blm-induced fibrosis. A) Schematic outlining tryptophan metabolism via the kyn pathway, processing steps are labeled as arrows, enzymes responsible for each step are shown in boxed italicized text. B) Mice were treated with 0.75U/Kg of bleomycin, or saline alone, and lungs were harvested at day 21. Lung lobes were flash frozen and prepared in a methanol extraction buffer for metabolomics analysis. C) Trichrome stain of histology of mice treated with saline, blm, or blm treated with 40 mg / kg kyn or kyn + 5 mg/kg CH223191, or vehicle by oral gavage on days 10-21, harvested on d21. Representative images of histology from 5 mice per condition. D, E, and F) Hydroxyproline assay on whole lungs of mice treated with saline, blm, or blm treated with 40 mg / kg kyn (panel F), 40 mg / kg of 3- hydroxykynurenine (3-OH-Kyn, panel G) or 80 mg / kg of 3-indole-propionic acid (3-IPA, panel H), or vehicle by oral gavage on days 10-21, harvested on d21. n=5-9 mice per condition. G) Experimental schematic showing days in which kyn was given via oral gavage. H) Total protein and relative IgM levels in mice treated with 40 mg / kg of Kyn or veh days 3- to 7-post blm treatment. I) Total protein of mice treated with veh or kyn in the fibrogenic period of blm recovery, days 10-21. n= 5 mice per group. Statistical significance was determined via ANOVA, * = p<0.05, **=p<0.01, ***=p<0.001.

Wang et al. recently reported that while kyn concentrations are increased in the lung following blm-induced fibrosis, kyn acts in an ameliorative or restorative fashion, reducing lung injury and fibrosis.(25) To further investigate the effects of kyn on both lung injury and fibrosis, we tested the effects of kyn at two time points: giving kyn for three days before instilling blm and continuing for seven more days until harvest at day 7 to test the role of kyn in the acute injury phase; or providing kyn to mice starting at day 10-post blm and harvesting at day 21 to test the effects of kyn during the fibrogenic period **(Figure 1G)**. We hypothesized that if kyn was increasing the tight junction function of epithelial cells in vivo, affecting migration of epithelial cells during wound healing, or diminishing inflammation, then kyn might be protective during the acute response to blm injury. However, if kyn was limiting the ability of epithelial cells to close a wound in the setting of fibrosis, then we would see more lung injury when kyn was administered during the fibrogenic period. Interestingly, although we observed evidence of lung injury in mice given bleomycin, as evidenced by increased total protein or IgM in BALF **(Figure 1H and 1I)**, There was no significant difference in lung injury between blm-treated mice given vehicle or kyn at either time point **(Figure 1H and 1I)**. We thus conclude that, contrary to what other groups have published, the effects of kyn are limited to fibrogenesis and do not alter early inflammatory events in the bleomycin model.

### Lung resident CD103+ DCs have complex biological functions that cannot be easily replicated in-vitro

Fibrotic lungs produce copious amounts of AHR activating tryptophan metabolites **(Figure 1)**. However, lung DCs do not undergo canonical AHR signaling even though they express the highest amounts of AHR in comparison to all other lung immune cells.(26) To further study the effects of kyn on lung DCs we isolated primary CD103+ DCs from the lungs of blm treated mice (7-days post blm) and grew them in standard DC cell culture conditions (complete RPMI media supplemented with 10% FBS). Complicating our initial study of these cells *ex-vivo*, lung DCs quicky took on an anti-inflammatory phenotype after just 12- to 18-hours in cell culture media with marked decrease in inflammatory transcripts, e.g. IL-6, and concomitant up-regulation of canonical AHR gene products like Cyp1b1, without the addition of exogenous AHR activating compounds **(Figure 2A)**.

**Figure 2.**
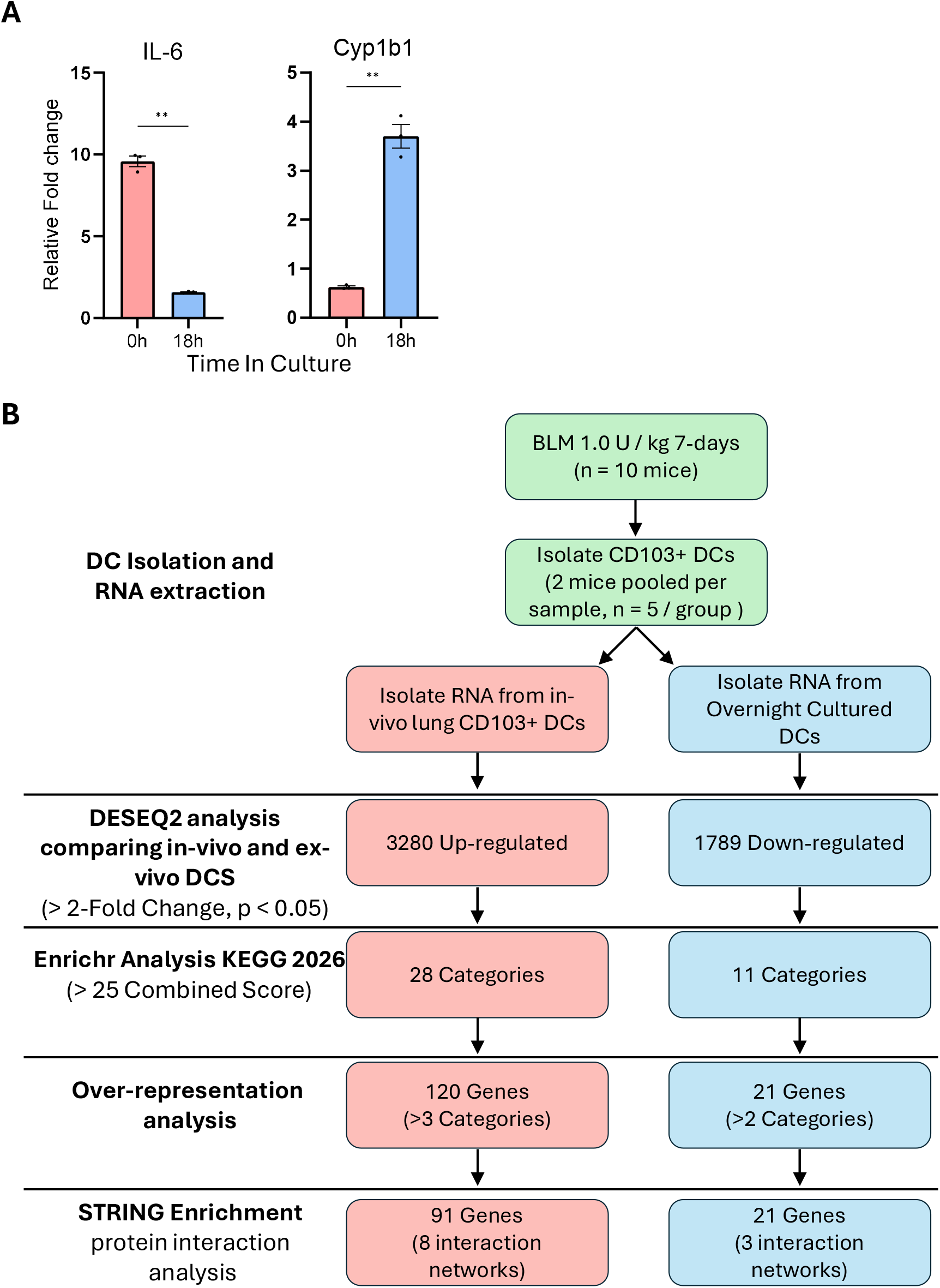
Bioinformatic pipeline for the comparative transcriptomic profiling of in vivo and in vitro dendritic cells. A) Mice (n = 5) were given 0.5 U / kg blm and lungs were harvested on d7. CD103+ DCs were purified from single cell suspensions of lung collagenase digestions. RNA was extracted either directly from lung DCs or from DCs cultured for 18h in RPMI culture media and analyzed for the indicated transcript by qPCR. B) High-throughput RNA-seq data from murine lung-derived (in vivo) and cultured (in vitro) dendritic cells were processed using DESeq2 to identify significantly differentially expressed protein-coding genes (p < 0.05, Foldchange > 2). Functional annotation profiles were generated via Enrichr analysis using a Combined Score >25. Over representation analysis identified recurring core driver genes appearing in multiple Enrichr categories. STRING analysis was conducted on identified core driver genes identifying networks of interacting genes in each condition.

To more fully understand the effects of cell culture conditions on lung DCs, we next investigated the effects of *ex-vivo* cell culture on lung CD103+ DCs by comparative transcriptomic analysis of freshly isolated DCs with DCs that had been cultured in media for 18h. Our magnetic bead isolation strategy using a combination of pan-DC isolation beads with a specific CD103+ bead purification yielded cells that were highly enriched for many lineage specific transcripts of conventional CD103+ DCs including xCR1, Clec9a, as well as Itgae (CD103) itself. Also noted was a cluster of highly expressed transcripts including various MHC genes as well as genes involved in antigen cross-presentation, i.e., Wdfy4, CD74, H2-Aa, H2-Ab1, and H2-Eb1 **(Supplemental Figure 1)**. These cells exhibited comparatively lower expression of genes associated with other leukocytes including cDC2, monocytes, macrophages, neutrophils, various lymphocytes, as well as fibroblasts and epithelial cells indicating that our cells were highly enriched for lung CD103+ DCs.

Bulk RNAseq and DEG analysis was conducted revealing 3280 up-regulated transcripts and 1789 down-regulated transcripts that were enriched in either tissue resident or cultured DCs respectively **(Figure 2B)**. To further investigate biological signatures of these cells, we conducted bioinformatic analysis using a combination of Enrichr and STRING analysis techniques to identify significantly enriched biological themes and statistically over-represented genes within those themes **(Figure 2B)**. Enrichr analysis identified 28 categories that were over- represented in the tissue DCs and 11 categories in the cultured DCs with a statistical combined score of > 25. The top two significantly enriched categories in the tissue resident DCs were extra-cellular matrix (ECM) interactions and integrin signaling **(Figure 3A)**. Tissue resident DCs were also enriched for many KEGG categories pertaining to cellular signaling pathways, like, Mapk, Rap1, and Pi3K-AKT signaling, as well as several categories pertaining to inflammation, i.e., cytokine receptor interactions and IL-17 signaling pathways **(Figure 3A)**. In contrast, cultured DCs were enriched for several KEGG categories pertaining to metabolism and activation of cytochrome P450 which fits with the profile of increased canonical AHR signaling that we observed.

**Figure 3.**
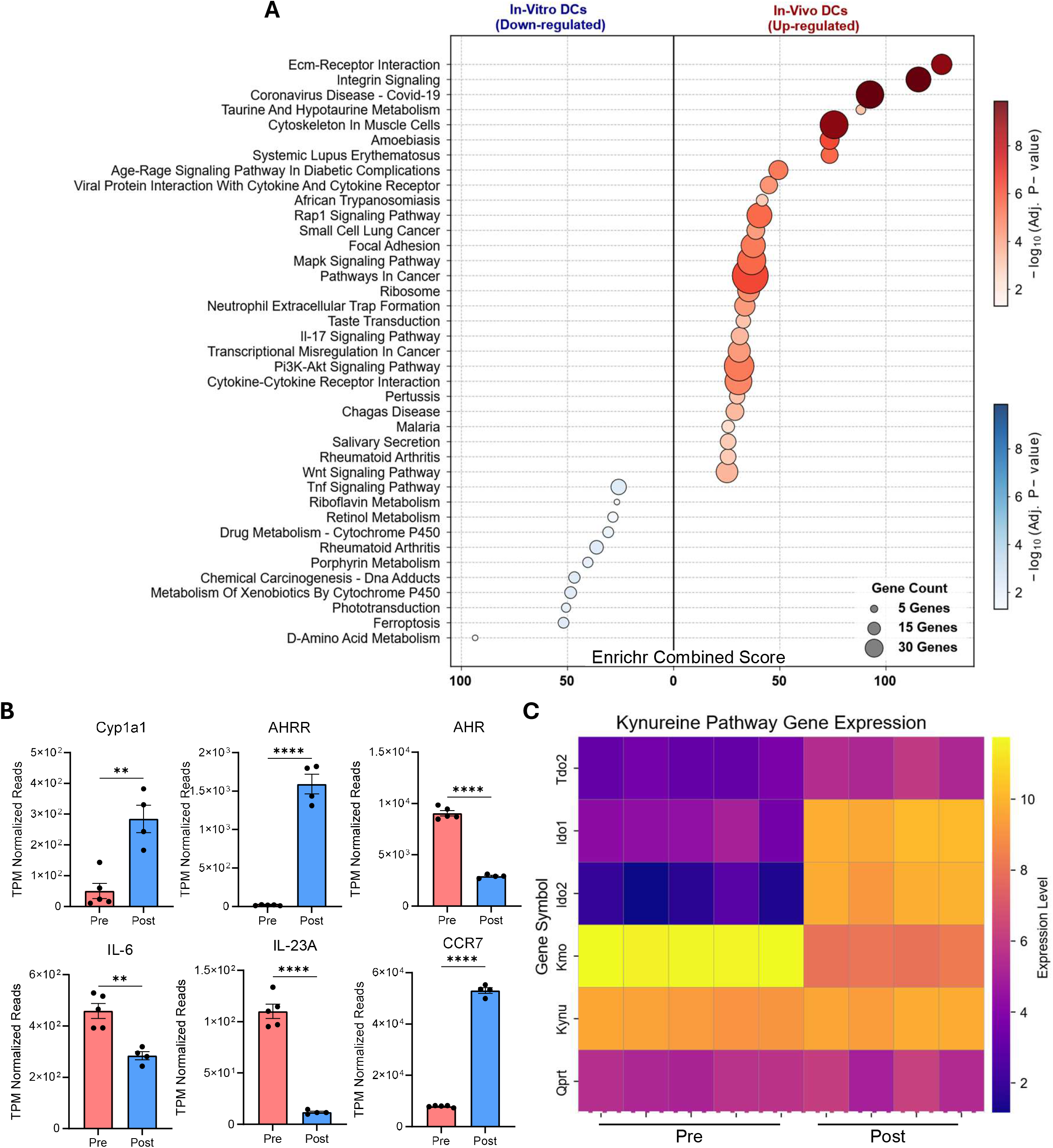
Enrichr analysis highlighting significantly enriched functional categories. A) Differentially expressed genes (fold change cutoff of > 2.0 and p < 0.05) were analyzed via Enrichr using the KEGG 2026 database. Categories with an Enrichr combined score of > 25 were picked as biologically meaningful. Categories enriched in tissue resident DCs (*In-vivo*) are shown in red while categories enriched in overnight cultured DCs (*ex-vivo*) are shown in blue. B and C) Select transcripts identified as significantly differentially expressed by DESEQ2 analysis.

Further analysis of individual transcripts confirmed robust activation of canonical AHR signaling in cultured DCs via up-regulation of the canonical AHR gene products Cyp1a1 and AHRR, however, expression of AHR itself was augmented in tissue DCs in comparison to cultured **(Figure 3B)**. Cultured DCs also expressed much lower IL-6 and IL-23A transcripts indicating that inflammation and induction of IL-17 pathways was diminished after 18h in cell culture **(Figure 3B)**. Interestingly, CCR7, a marker of DC maturation was repressed *invivo,* indicating that the fibrotic lung milieu may inhibit some aspects of maturation and keep immature DCs from migrating to lung draining lymph nodes **(Figure 3B)**. Mature DCs are well known to produce the tryptophan processing enzyme IDO1.(41) We thus analyzed our DCs for expression of IDO1, IDO2, and TDO2 as well as other enzymes involved in processing tryptophan along the KP. Interestingly, tissue resident CD103+ DCs exhibited very low expression of IDO1 and IDO2 which was dramatically up-regulated after cell culture **(Figure 3C)**. Kynurenine mono-oxygenase (KMO), the enzyme responsible for further processing kyn to HK, showed the opposite phenotype and was highly up-regulated in tissue resident DCs in comparison to cultured DCs **(Figure 3C)**. Minimal change in expression was observed for the enzymes TDO2, KynU, and QPRT, indicating that tissue resident CD103+ DCs express a unique profile of tryptophan processing enzymes that is changed when they are cultured *ex vivo*.

To further process gene signatures from the Enrichr analysis, we utilized an over-representation analysis technique that identified core driver genes that appeared in multiple Enrichr categories simultaneously. We identified 91 core driver genes in tissue resident DCs and 21 core drivers from cultured DCs **(Figure 2B)**. We utilized the STRING analysis package within cytoscape in conjunction with MCL clustering to group these core driver genes into interacting networks and identified 8 networks in tissue DCs and 3 networks in cultured DCs **(Figure 4)**. Three large interacting networks were identified in tissue DCs that had biological themes revolving around TLR / NfκB and IL-17 signaling, ECM interacting proteins, and growth factors. We also identified five smaller interacting networks that had genes related to lipid processing, cell cycle, complement, ephrin signaling, and type IV collagens **(Figure 4)**. Cultured DCs had three interacting networks, the largest of which had genes involved in canonical AHR signaling, while the two smaller networks dealt with cytokines / chemokines and ferroptosis **(Figure 4)**. We also processed these lists of core driver genes with GSEA, looking for any enriched transcription factors that might drive these biological phenotypes. Interestingly, only one transcription factor was significantly enriched, RelB (p = 0.03, gene symbols denoted with red asterisks in **Figure 4**). RelB is a known dimerization partner with AHR and is important in driving non-canonical inflammatory AHR signaling.(24, 42) In summation, Enricher analysis indicates that lung resident CD103+ DCs are undergoing complex cell-cell interactions in the fibrotic lung and are participating in an inflammatory IL-17 centric circuit via non-canonical AHR signaling, which agrees with the role we had previously described for these DCs in both bleomycin -induced fibrosis and IPF.(43)

**Figure 4.**
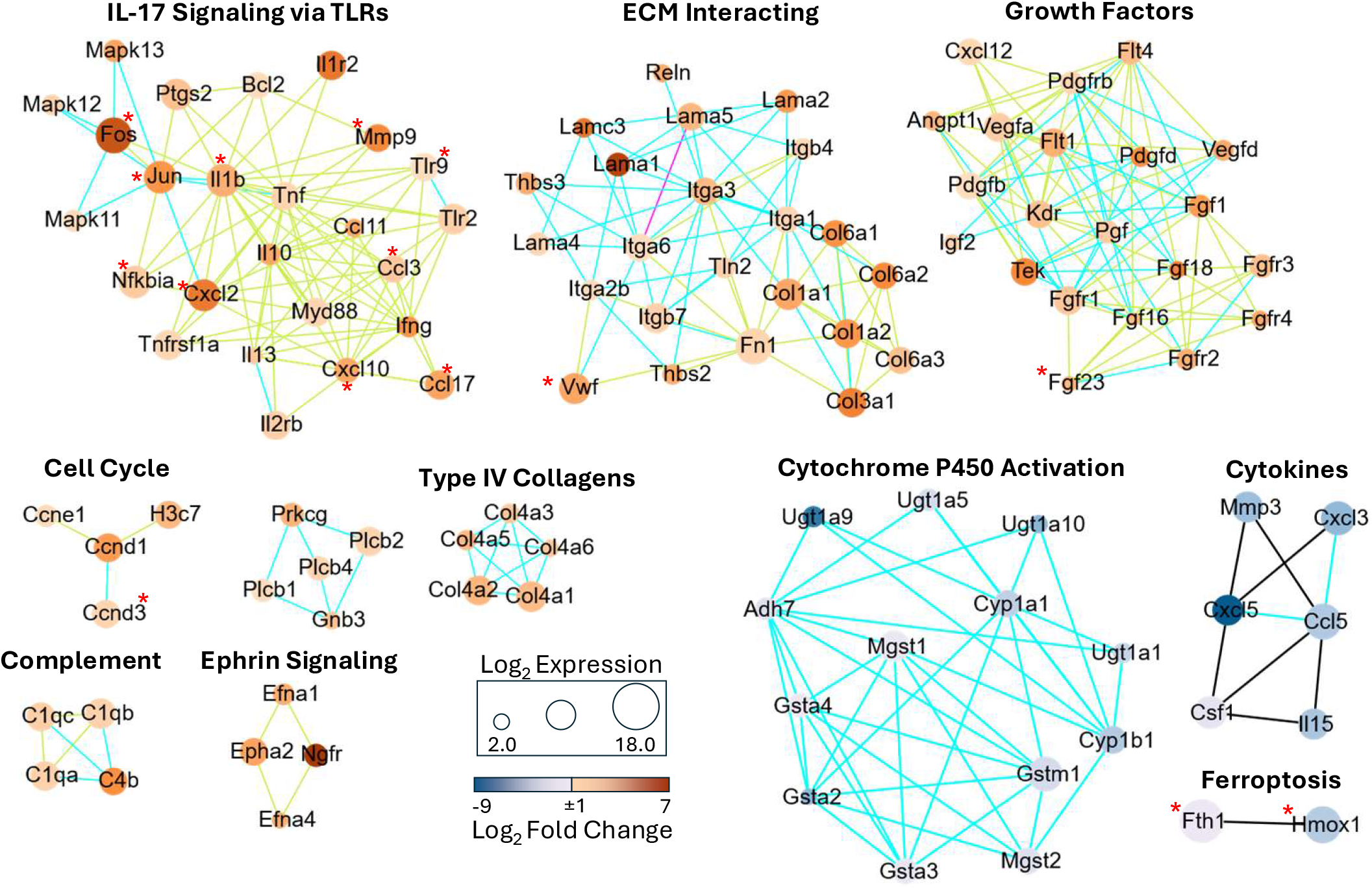
Identification of functional gene networks in in vivo and ex vivo DCs. STRING analysis was performed in Cytoscape 3.10.4 to identify interactions of genes that appeared in > 3 functional categories (up-regulated genes, in-vivo DCs) or > 2 functional categories (down- regulated genes ex-vivo DCs). MCL clustering was used (inflation value 2.5) to identify sub- networks of interacting genes. Circle size denotes the overall expression of the transcript (TPM normalized reads), circle color denotes overall fold-change. Edge color is reflective of the STRING interaction type: blue = database curated, green = text mined, black = co-expression, magenta = experimentally validated, red asterisks next to gene symbols indicate RelB target genes identified via GSEA.

To further probe the effects of kyn on DCs, we generated inducible CD103 DCs (iCD103) or purified lung CD103+ DCs and cultured them in the presence of the TLR9 ligand ODN2395 with or without the addition of the tryptophan metabolites kyn or 3OH-kyn. We chose ODN2395 based on our previous study that identified an important role for TLR9 in bleomycin induced fibrosis(26) which was further supported by our RNAseq of tissue resident CD103+ DCs that identified TLR9 as significantly up-regulated **(Figure 4)**. While stimulation with ODN2395 drove strong expression of IL-6 in both iCD103 cells as well as tissue isolated CD103+ DCs, co-treatment of DCs with kyn or HK strongly repressed IL-6 expression and instead drove expression of the canonical AHR gene product Cyp1b1 **(Figure 5A and 5B**). We next attempted to further re-capitulate the microenvironment of the fibrotic lung by treating iCD103 DCs with ODN2395 in conjunction with either hypoxia or TGF-β. AHR and the hypoxia-inducible pathway have long been considered antagonistic due to the shared binding partner ARNT, also known as HIF1-β.(44) Hypoxia (2% O_2_), however, did not modulate the effect of AHR on the inflammatory or canonical pathway **(Figure 5C)**. TGF-β is often anti-inflammatory and has been shown to antagonize AHR in other systems(45, 46), however TGF-β increased both canonical AHR response and IL-6 response while not affecting AHR levels **(Figure 5D)**, also failing to recapitulate in vivo CD103+ phenotypes.

**Figure 5.**
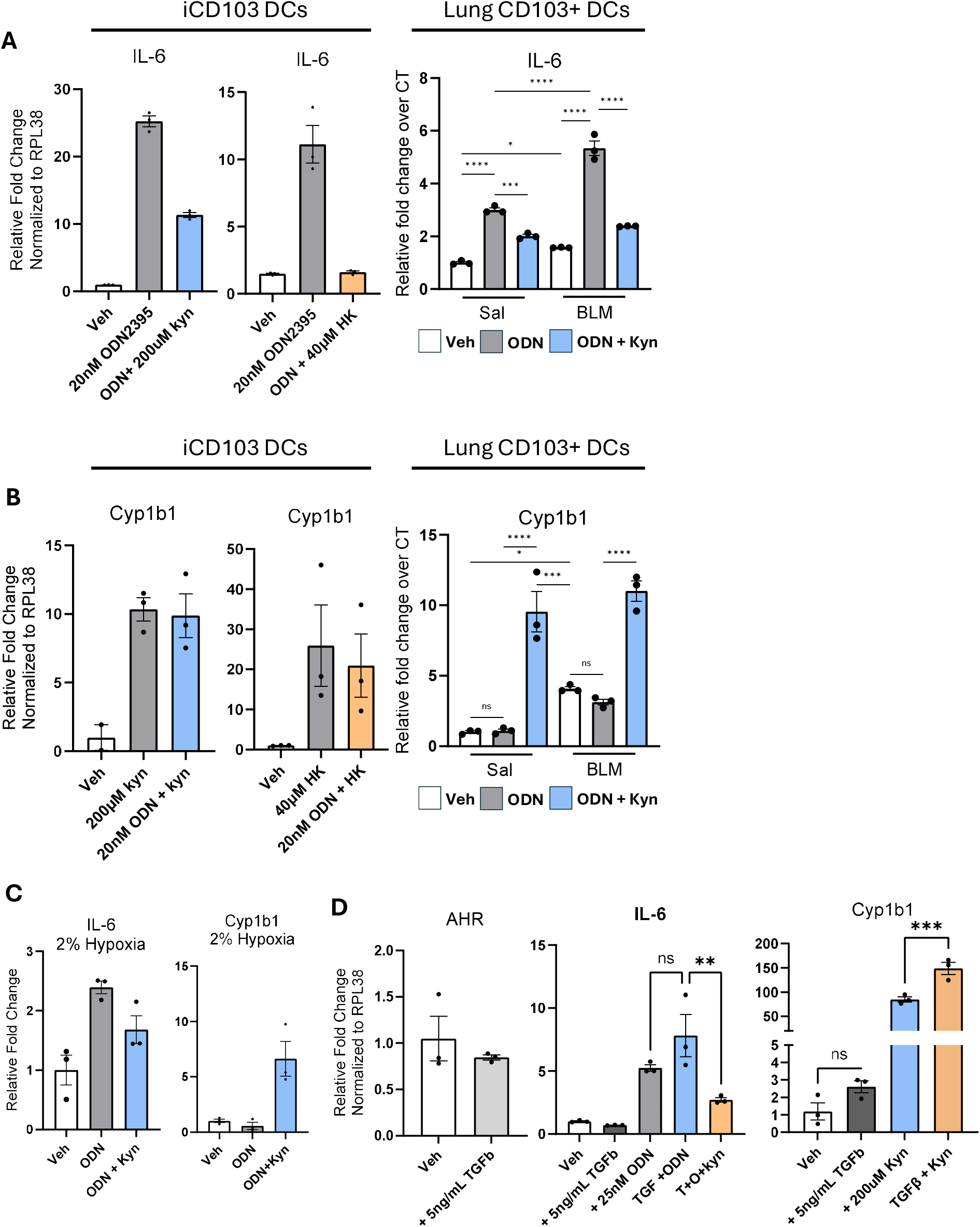
*Ex vivo* culture of DCs does not recapitulate the *in vivo* phenotype. (A-B) qPCR results of IL-6, Cyp1B, and AHR from DCs treated overnight with the described conditions. C) Media treatments were consistent as described in A, and cells were housed overnight in 2% O_2_ to induce hypoxia. D) Cells were treated overnight with TGFβ (5 ng / mL) and described conditions. Significance determined by one-way ANOVA. Representative of at least two independent experiments. Statistical significance was determined by ANOVA, * = p < 0.05, ** = p < 0.01, *** = p<0.001, **** = p < 0.0001.

### CD103+ dendritic cells are situated in fibrotic foci and co-culture of DCs with fibroblasts reverses the anti-inflammatory effects of kyn

Given that *ex vivo* mono-culture of lung DCs did not recapitulate the *in vivo* inflammatory phenotypes of the fibrotic lung, we aimed to analyze mechanisms by which DCs are contributing to fibrosis. We first evaluated if these scarce cells were present in fibrotic foci via immuno-fluorescent histochemistry. We administered blm to GFP-Col mice in which all collagen producing cells (mainly fibroblasts) are tagged with GFP and found that CD11c / CD103 double positive cells, representing conventional type 1 DCs, were present in fibrotic foci specifically near highly activated GFP+ fibroblasts **(Figure 6A)**. Interestingly, CD11c / CD103 double positive cells were exceedingly rare in areas of normal lung architecture but concentrated in pockets of remodeled fibrotic tissue **(Figure 6B)**. Flow cytometry of immune cells isolated from saline or blm-treated lungs treated with kyn or a vehicle control from days 10-14 post-blm did not show any difference in immune populations by day 14 **(Supplemental Figure 2A)**. However, administration of kyn to control saline treated mice did upregulate expression of the canonical AHR gene product Cyp1b1, while blm-treated mice failed to upregulate cyp1b1 when given the same dose of kyn **(Supplemental figure 2B)**. Tissue resident DCs express many genes involved in sensing and interacting with the complex ECM that is present during fibrosis **(Figure 3 and Figure 4)**. Given that we suspect DCs are directly associated with fibroblasts in the fibrotic lung, we next employed a co-culture strategy wherein we harvested primary murine lung fibroblasts from mice at 21-d post -blm instillation and directly cultured these fibroblasts with ex-vivo generated iCD103 cells. As shown earlier, mono-cultured iCD103 DCs suppressed production of IL-6 when exposed to kyn in the cell culture media **(Figure 6C)**. Upon co-culture with lung fibroblasts, the exact opposite phenotype was observed in that kyn augmented production of IL-6 **(Figure 6C)**. Additionally, kyn administration uniquely drove production of Cyp1a1 in mono-cultured DCs, but activation of canonical AHR was suppressed when DCs were co-cultured with fibroblasts **(Figure 6D)**. To test whether this phenotype was driven by soluble factors or by cell-cell contact, we cultured iCD103+ cells with BALF harvested from mice 21-d post blm. BALF treatment effectively lowered expression of Cyp1A1, indicating decreased canonical AHR signaling, however, no change was observed in production of IL-6 **(Figure 6E)**. These data indicate that a soluble factor produced in the fibrotic lung is sufficient to alter AHR signaling but was insufficient to drive inflammation.

**Figure 6.**
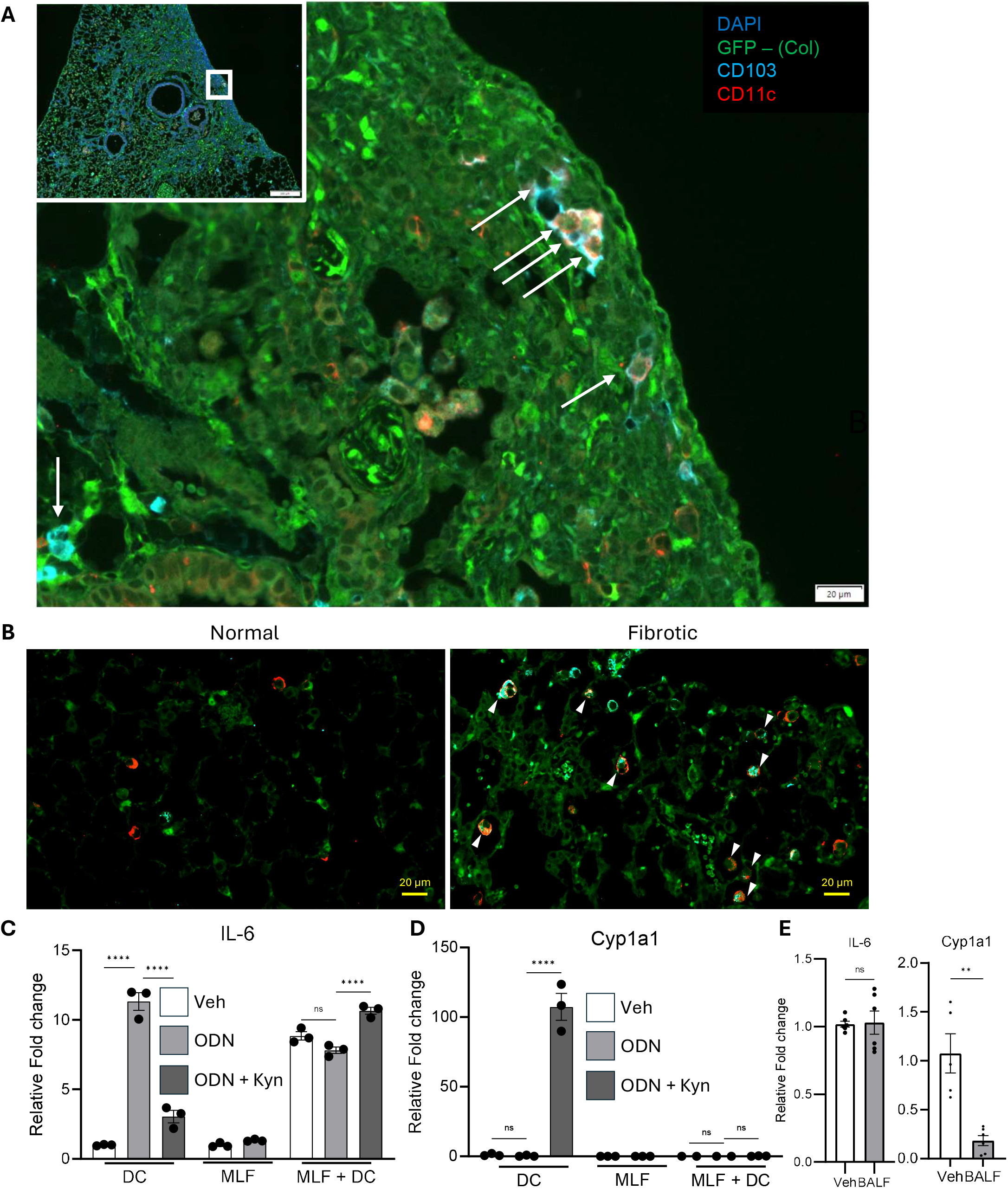
CD103 + cDC1s exist in fibrotic foci and exhibit inflammatory phentoypes when co-cultured with structural cells *ex-vivo.* A) IHC staining showing CD103+/CD11c+ cells in fibrotic foci. GFPCol1a1 mice were given fibrosis via blm instillation, and lungs were fixed in 10% buffered PFA on day 21. Lungs were processed and stained for IHC against GFP, CD103, and CD11c. Arrows indicate cells with red/cyan staining. Inset: wide view. B) Additional images showing the presence of CD103+ CD11c+ DCs in fibrotic but not normal areas of lung sections obtained from mice with bleomycin induced fibrosis. C and D) iCD103 DCs were mono-cultured in complete media with 25 nM ODN2395 (ODN) or ODN + 100 μM kyn (ODN + Kyn), or co- cultured with 1.5×10^5^ primary murine lung fibroblasts (MLF + DC) isolated from lung minces of mice with bleomycin induced fibrosis (21-days post bleo), MLF cells were cultured alone as control. n = 3 per group. E) iCD103 DCs were incubated with 50% BALF from mice 21-d post blm. RNA was isolated and analyzed for the indicated transcript (n = 5-6 samples per group. Statisitcal significance was assessed via ANOVA, ** = p < 0.01, **** = p < 0.001.

### Fibroblast activation is not affected by AHR signaling by kyn

AHR has previously been reported to affect myofibroblast activation.(44, 47–49) However, these studies did not use lung specific AHR ligands, i.e., tryptophan metabolites. To study the effects of lung relevant AHR ligands on fibroblast activation, we used the endogenous ligand kyn in conjunction with the AHR inhibitor CH223191. CCL210 cells, a human fibroblast cell line, were activated with TGF-β to induce myofibroblast differentiation and subsequently treated with kyn or CH223191. As expected, TGF- β increased the expression of α-smooth muscle actin (αSMA) **(Figure 7A)**. However, AHR activation had no effect on αSMA expression **(Figure 7A quantified in 7B)**. Similarly, there was no significant difference in *Col1a1*, α*SMA,* or *AHR* transcript expression as measured by qPCR, and TGF-β did not affect the expression of *Cyp1b1* induced by kyn in these cells **(Figure 7C)**. We further tested the effects of AHR activation in primary murine lung fibroblasts isolated from either saline or blm treated mice. Interestingly, kyn treatment slightly decreased migration in fibroblasts isolated from saline control mice as assessed by scratch wound assay, however, no significant effect was observed in fibroblasts isolated from blm mice **(Figure 7D)**. Inhibition of AHR with CH223191 did not significantly affect the mobility of either group of fibroblasts by scratch wound assay **(Figure 7E)**.

**Figure 7.**
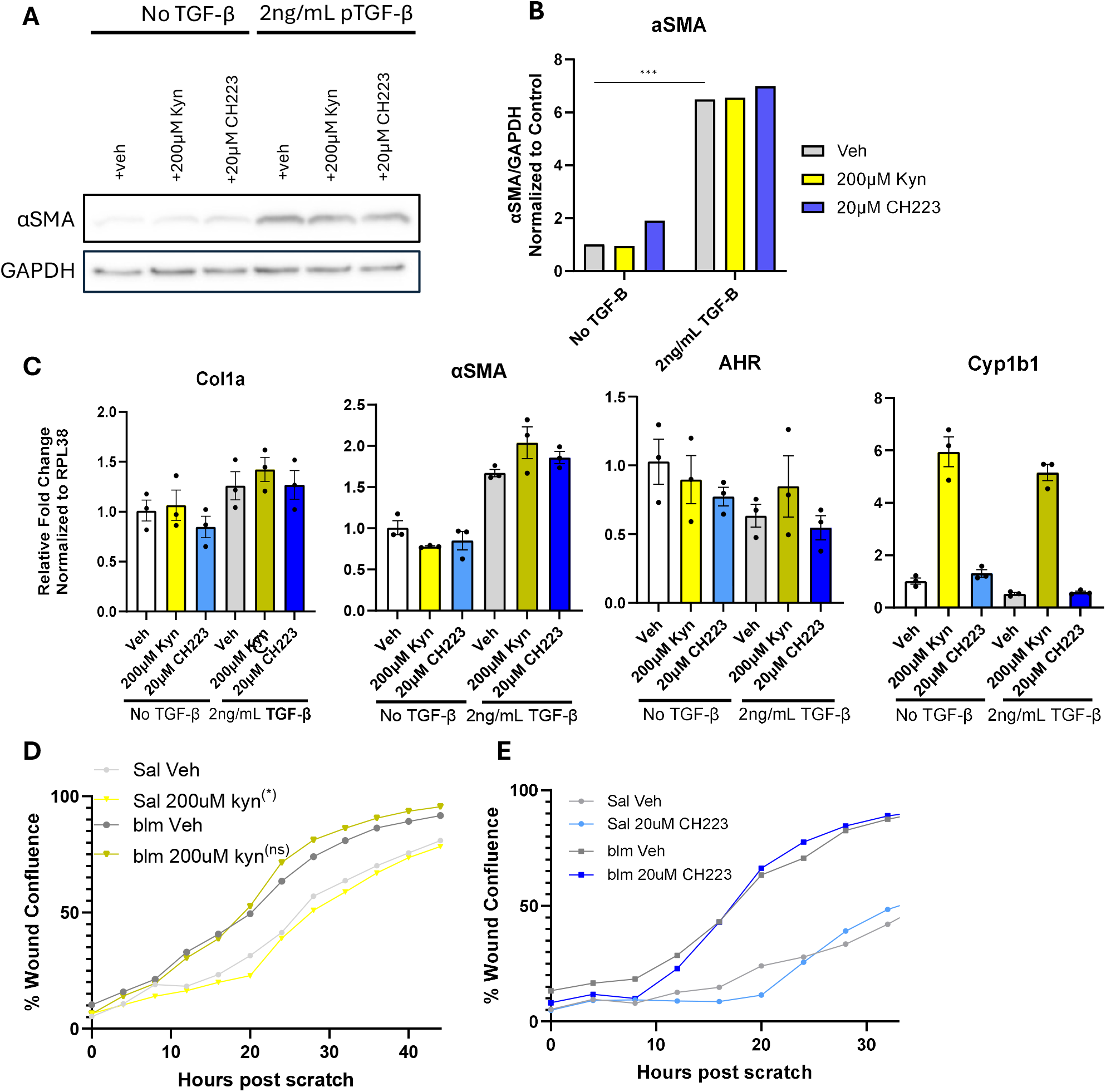
Fibroblast activation is not altered by kyn or CH223191 treatment. A) CCL210 fibroblasts were plated at 4×10^5^ cells per well of a 6-well plate. Cells were treated with veh, 2ng/mL TGF-β, 200uM kyn, and/or 20uM CH223191 as noted. B) Western blot was quantified using ImageJ. C) qPCR results of Col1a1, Cyp1B, αSMA, and AHR RNA from CCL210 cells treated with TGF-β, then kyn or CH223191 as noted. D) Fibroblasts isolated from mice at 21d post saline or blm instillation were isolated, plated to 90% confluence, and treated with veh, 200uM Kyn, or 20uM CH223191, then scratched with a Sartorius Woundmaker, and imaged consecutively in an Incucyte before being analyzed by Sartorius Incucyte software. All data representative of at least two independent experiments.

### In epithelial cells, TGF-β boosts AHR signaling, which affects epithelial cell migration

Zimmerman et al. recently described a role for AHR in modulating both inflammatory phenotypes as well as tight barrier functions in pulmonary epithelial cells.(36) However, this study focused on FICZ, an AHR ligand with an affinity thousands of fold higher than that of kyn and other tryptophan metabolites. To assess the role of tryptophan metabolites and AHR activation on epithelial wound healing responses, we utilized MLE-12 cells, a bronchoalveolar epithelial cell line, using a scratch wound assay. We found that kyn alone was not sufficient to change the rate of wound closure, however, addition of 5 ng / mL TGF-β alone modestly increased the rate of wound closure **(Supplemental Figure 3A)**. When TGF-β was added in conjunction with 200µm kyn, we observed the opposite phenotype in that MLE-12s were impaired in wound closure **(Supplemental Figure 3A)**. We next tested whether FICZ would alter epithelial cell migration. In agreement with Zimmerman et al., FICZ reduced migration, however, FICZ did not require the AHR amplifying effects of TGF- β to achieve this **(Supplemental Figure 3A)**. To further assess the effects of TGF- β on both AHR expression and signaling we treated MLE-12 cells with TGF-β and analyzed expression of AHR itself as well as the canonical AHR gene product Cyp1b1 by qPCR. Treatment with TGF-β increased expression of *AHR* and *Cyp1b1* in response to kyn, which could be inhibited by addition of CH223191 **(Supplemental Figure 3B-C)**. Thus, we conclude that the effects of TGF-β amplify AHR expression and signaling allowing kyn to reduce epithelial cell migration, albeit modestly.

To address if kyn would have the same effect on primary lung cells, we isolated AECs from saline or blm-treated mice and cultured these cells with additional kyn. AECs from fibrotic mice showed a trend towards increased expression of *Cyp1b1* at baseline but did express more when treated with kyn **(Supplemental Figure 3E).** To address if this effect was AHR specific, cells were co-treated with the AHR antagonist CH223191, which ameliorated *Cyp1b1* expression in both saline and blm AECs **(Supplemental Figure 3E)**. To address if kyn would impact tight barrier function we grew AECs to confluence under an air liquid interface and simulated an inflammatory “wound” with the addition of TNFα and IFNγ with or without the addition of kyn. Although the wounding process did reduce tight barrier function as shown by an initial decrease in TEER. We did not see any difference in TEER response to injury when 200µM kyn alone was added to the primary AECs **(Supplemental Figure 3F)**. Thus, we conclude that while strong AHR agonists, like FICZ, can have effects on pulmonary epithelial cells, weaker AHR ligands require additional amplifying signals to achieve biological effects.

## Discussion

### Tryptophan Metabolomics in the blm-induced fibrotic lung

It has long been appreciated that the kyn : tryptophan ratio is increased in the serum and lung tissue of IPF patients as well as in blm-treated mice. (25, 38, 50) However, previous reports ignored other tryptophan metabolites along the kynurenine, serotonin, or bacterially-derived pathways. We found that there were significant differences in only a few metabolites, including tryptophan itself, kyn, HK, and IPA **(Figure 1B)**. This suggests that increased tryptophan metabolites are pooling early in the KP and are not being fully metabolized to NAD+ and are thus independent of energy production. We have previously shown that kyn worsens fibrosis when delivered during the fibrogenic period of the blm mouse model, however, HK and IPA had not been tested. Although both compounds are AHR agonists **(Figure 5A and ref** (51)), HK only showed a trend towards increased fibrosis that did not reach statistical significance, and IPA had no effect **(Figure 1D, 1E, and 1F)**. Therefore, we continued to focus primarily on kyn and the AHR antagonist CH223191 in subsequent experiments.

### In vitro conditions do not replicate in vivo phenotypes in monoculture

Our previous work identified that dendritic cells (DCs) in the blm-induced fibrosis model responded to kyn by undergoing non-canonical AHR signaling to promote fibrogenesis *in vivo*. Thus, we were eager to identify the reasons that DCs preferentially signaled via the non-canonical AHR pathway, and to do so, wanted to culture the cells *ex vivo*. However, within 18 h of being out of the fibrotic lung milieu, the *ex vivo* DCs preferentially signaled via canonical AHR, evidenced by up-regulation of Cyp1b1 **(Figure 2A)**. Canonical AHR is generally immunosuppressive partly due to activation of NF-κB suppressing genes such as TiPARP.(52, 53) Cultured DCs quickly down-regulated expression of inflammatory gene products which was seen in the transcriptomes of the freshly isolated vs. cultured DCs (**Figure 2A, 3, and 4**). We next tried to identify relevant conditions to culture the DCs *ex vivo* in an effort to recapitulate the *in vivo* phenotype. Initially, we tested the effects of high concentrations (200µM) of kyn in the context of a TLR9 agonist, ODN2395, which simulates inflammation from a sterile trigger. We found that both kyn and HK, the immediate metabolite of kyn, activate canonical AHR and reduce inflammation *ex vivo* in both bone marrow generated iCD103 DCs as well as lung CD103+ DCs isolated from either saline or blm challenged mice **(Figure 5A-B)**.

Similarly, hypoxia has been linked to canonical AHR antagonism, but in our hands did not alter the response to kyn, reducing inflammation and increasing canonical signaling **(Figure 5C)**. We further introduced TGF-β, a cytokine which drives fibrosis and is highly expressed during wound healing. Interestingly, found that *ex vivo* administration TGF-β amplified IL-6 production in response to ODN stimulation, however, the addition of kyn in combiniation with TGF-β was immunosuppressive and actually amplified production of canonical AHR responses **(Figure 5D)**. Other mechanisms that might account for altered AHR signaling including tryptophan concentrations (54), cAMP concentrations (55), cell-to-cell contact (56), and DC maturation state.(57, 58) Given that CD103+ DCs were found in close approximation to fibroblasts in vivo (**Figure 6A**), and our previous work had identified fibroblast expression of TDO2, an enzyme capable of generating kyn, to be important in *vivo*, we opted next to co-culture the DCs with fibroblasts.

### Co-culture of CD103+ DCs with fibroblasts recapitulates non-canonical AHR signaling

When DCs were co-cultured in the presence of fibrotic fibroblasts, kyn stimulation now resulted in proinflammatory IL-6 production without induction of canonical AHR targets **(Figure 6C, D)**, and this was from the DCs, not the fibroblasts alone. In fact, fibroblast co-culture induced proinflammatory IL-6 from DCs even in the absence of ODN stimulation. To determine whether a fibrotic cytokine milieu might be sufficient to switch DC signaling from canonical to non-canonical AHR signaling, we tested monoculture of the DCs with BALF obtained from bleomycin-treated mice. While the BALF did inhibit canonical AHR signaling as measured by Cyp1a1 expression, it was insufficient to upregulate IL-6. Thus, we conclude that the *in vivo* phenotype which results in non-canonical AHR signaling in DCs in the fibrotic lung likely requires cell-cell contact with fibrotic fibroblasts. We next wanted to explore the impact of kyn treatment on fibroblasts in monoculture.

### Fibroblast activation is not significantly affected by kyn treatment

Fibroblasts play the most direct role in fibrogenesis, activating to become myofibroblasts, which deposit an excess of extracellular matrix. Previous studies have found that AHR activation blocks the activation of fibroblasts and protects from fibrosis (25, 48, 59, 60). However, this was opposite of our *in -vivo* findings. These previous studies have largely used strong AHR ligands exogenous to natural systems. We therefore evaluated how treatment with kyn and CH223191 affected αSMA production and expression of *Col1a1* and *Act2*. We tested CCL-210, a human lung fibroblast cell line, and found that neither the baseline cells nor the cells activated with TGF-β showed any significant change in αSMA protein expression or *Col1a1* or *Act2* RNA expression when treated with kyn or CH223191 **(Figure 7A-B)**. AHR expression itself was not altered by these conditions, although canonical signaling was induced by kyn treatment **(Figure 7C)**.

Using primary lung fibroblasts isolated via the crawl-out method from mice treated with saline or blm, we saw that fibroblasts from blm-treated mice had increased migratory capacity to close a scratch wound relative to fibroblasts from saline-treated mice. Kyn slightly increased the rate of wound closure in saline treated fibroblasts, however, no effect of kyn or CH223191 was noted in blm treated fibroblasts **(Figure 7D)**,and the overall effect of kyn, even in the saline fibroblasts, was minimal. We conclude that AHR activation via kyn or inhibition via CH223191 ex vivo did not alter the ECM deposition or the migratory capacity of fibroblasts. These results are also at odds with our finding that kyn increases deposition of collagen post-bleomycin in vivo, **(Figure 1)**, suggesting that *ex vivo* monoculture of fibroblasts cannot accurately capture the *in vivo* AHR signaling patterns or that the in vivo phenotype relies heavily on the DC output.

Our past studies have shown that fibroblasts play an important role in generating the kyn signal because they are the only cell type in the lung to express TDO2, and a TDO2 inhibitor protects from blm-induced fibrosis. This result was surprising to us given that tryptophan can also be converted to kyn via the IDO1 enzyme, which is more ubiquitously expressed. Other studies have indicated a potential role for IDO1 in pulmonary fibrosis.(61) However, our previous study did not replicate these data and indicated low overall expression of IDO1 in the lungs of IPF patients and in blm induced fibrosis, furthermore, IDO1 knock out mice showed no abrogation of fibrogenesis *in vivo* (26). Interestingly, tissue resident CD103+ DCs do not express IDO1 or IDO2 but do express elevated levels of the KP enzyme KMO **(Figure 3C)**, indicating that lung CD103+ DCs do not directly play a role in processing tryptophan but may further process kyn to HK *in vivo*. Importantly, our IHC studies do show the close association of CD103+ DCs with fibroblasts *in-vivo* **(Figure 6A and 6B)**. Together, these data suggest that proximity of DCs to fibroblasts may facilitate kyn-AHR signaling and promote fibrogenesis and further supports the hypothesis that TDO2 rather than IDO1 is important for fibrogenesis *in vivo*.

### TGF-β boosts AHR activity, but kyn does not affect function of epithelial cells

Epithelial cells are arguably the inciting player in pulmonary fibrosis, as their dysregulated wound healing may perpetuate fibrosis (62). We initially studied epithelial cells using MLE-12 cells, a cell line derived from mouse lung epithelial cells. In contrast to previous reports, which have found that AHR and TGF-β are antagonistic (46, 63–66), we found that treatment of MLE-12 cells with TGF-β boosted AHR expression and downstream signaling **(Supplemental Figure 3A-3C)**. Accordingly, while kyn alone was insufficient to change the wound-healing activity of MLE-12s from baseline, the addition of TGF-β was sufficient to induce AHR to decrease wound closure, albeit modestly **(Supplemental Figure 3A)**. AECs from blm-treated mice were also more primed to respond to kyn and produce more *Cyp1b1* than AECs from healthy mice **(Supplemental Figure 3E)**. While we did not see any direct ability of kyn to influence TEER in primary cells, as has been previously reported with the strong AHR agonist FICZ (36), we were unable to test the hypothesis that TGF-β would amplify the effects of kyn as TGF-β independently regulates TEER (67, 68).

Overall, our results show kyn can modestly reduce migration when TGF-β is present suggesting that kyn may regulate wound healing of epithelial cells in the fibrotic lung. To determine the epithelial impact of kyn and AHR signaling *in vivo*, we measured lung injury by total protein and IgM concentration in bronchoalveolar lavage fluid (BALF) of blm mice treated daily with kyn either early (days -3 to 7-post blm) or late (days 10- to 21-post blm). Previous research has shown that AHR signaling bolsters epithelial barrier function and controls cell response to damage (36, 69, 70). These studies suggest that kyn treatment during the acute, inflammatory period of blm treatment would be protective. However, kyn treatment before and for a week following blm did not result in a change in lung leak at one week post blm. Similarly, kyn treatment during the fibrogenic periods did not result in increased lung leak at day 21 **(Figure 1G and 1H)**. Overall, this suggests that kyn treatment is insufficient to significantly alter epithelial function *in vivo* in the lung and shows that in AECs, *ex-vivo* conditions do not mirror AHR effects *in-vivo*.

### AHR as an inflammatory mediator in vivo

In our previous paper, we focused on the importance of dendritic cell AHR signaling in pulmonary fibrosis, where we showed that pharmacologic inhibition and genetic AHR manipulation in DCs could modulate fibrotic outcomes.(26) Dendritic cells make up a very small number of cells in the lung but are increased in blm (71) and found in close proximity to activated, collagen producing, fibroblasts in fibrotic foci **(Figure 6A and 6B)**. This is consistent with our previous results, which identified fibroblasts as the source of TDO2-derived kyn, which activated the DCs.(26) While activation of AHR is known to affect the maturation and migration of various immune cells, including dendritic cells and monocytes (72–74), We did not find that treatment with kyn affected immune cell populations during blm induced fibrosis, leading us to conclude that AHR modulation in the fibrotic lung solely influences cellular function without affecting leukocyte recruitment **(Supplemental Fig. 2)**.

As discussed above, monoculture of either primary lung or iCD103 DCs with kyn exhibited an anti-inflammatory effect driven by induction of canonical AHR signaling. Our previous work showed that CD103+ DCs directly isolated from fibrotic lungs exhibited no Cyp1b1 expression, low TIPARP, and high IL-6 expression even though fibrotic lungs produce large quantities of kyn. Furthermore, the hyperinflammation was AHR dependent, as production of IL-6 decreased when AHR was antagonized.(26) Importantly, co-culture of DCs with primary lung fibroblasts reversed the effects of kyn, allowing transcription of inflammatory IL-6 while suppressing activation of canonical AHR signaling **(Figure 6C and 6D)**.

There are numerous possible explanations for why *ex vivo* culture cannot replicate *in vivo* phenotypes. Given that co-culture of DCs with lung fibroblasts reverses the anti-inflammatory effects of kyn it is likely that cell-to-cell or cell-to-matrix contact is necessary for AHR-dependent inflammation. This hypothesis is reinforced by the myriad ECM interacting genes we identified as being up-regulated in lung DCs that was lost upon *ex vivo* culturing **(Figure 4)**. ECM interacting proteins, especially integrins, are extremely important in driving fibrosis and interact in both sensing the altered ECM present in fibrotic tissue as well as processing key fibrotic cytokines such as TGFβ.(75–77) While little is known specifically about how DCs utilize ECM sensing integrins to modulate fibrotic signaling, other studies have identified interactions between various integrins and AHR, which drives tissue invasion of pathogenic cells and TGFβ processing. (78, 79). It is important to understand the cellular context of AHR signaling as new AHR modulating therapies are coming to market. The newly FDA approved drug Tapinarof treats psoriasis through AHR agonism, opening the door for potential treatment of other diseases in which AHR is implicated. It is important, therefore, to consider how different cell types and disease states may affect AHR signaling. We demonstrate here how AHR signaling *in vitro* is not necessarily relevant to *in vivo* function in monoculture, and therefore future studies should consider the limitations of *in vitro* findings for AHR regulation of primary lung cells and consider important cell-cell interactions. Our results also suggest that *in vivo* agonism of AHR could promote fibrogenesis, which is a cautionary tale for careful monitoring of patients on AHR modulating drugs, like Tapinarof.

## Supporting information

Table of PCR primers

## Author Contributions

HC conceived and designed research, performed experiments, analyzed data, interpreted results, prepared figures, and drafted manuscript. RM performed experiments, analyzed data, and prepared figures. BA performed experiments, analyzed data, interpreted results, and edited manuscript. JF and JK performed experiments and analyzed data. KJ performed experiments, analyzed data, and interpreted results. RZ designed experiments and prepared figures. BM designed experiments, interpreted results of experiment, edited and revised manuscript, and approved final version of manuscript. SG designed and performed experiments, analyzed data, interpreted results, prepared figures, wrote, edited and revised manuscript, and approved final version of manuscript.

## Data Availability

Raw and processed RNAseq datasets are made available on the Gene Expression Omnibus database (GSE338791).

## Acknowledgments

Metabolomics measurements were performed by the University of Michigan Medical School BRCF Metabolomics Core Facility (RRID:SCR_026721). Ohmmeter was generously loaned by Tom Schmidt and Nicole Cady at the University of Michigan. The authors would like to thank the University of Michigan Flow Cytometry Core and the contributions of Sean Linkes and Kamlai Saiya-Cork for their contributions in processing and analysis of samples.

## Figure Legends

**Supplemental Figure 1.**
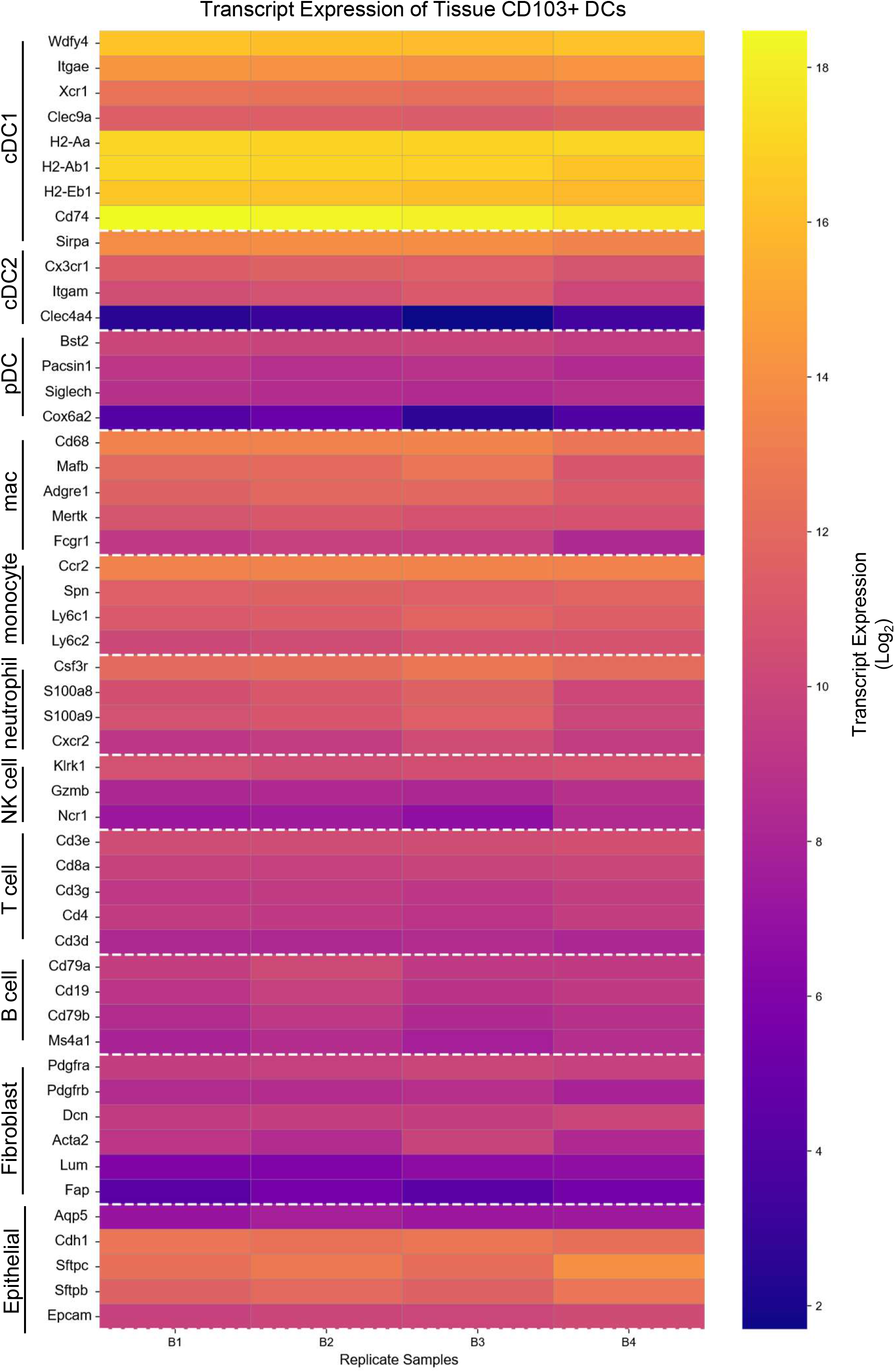
Heatmap showing expression of lineage specific transcripts taken from bulk RNAseq of bead purified lung CD103+ DCs.

**Supplemental Figure 2.**
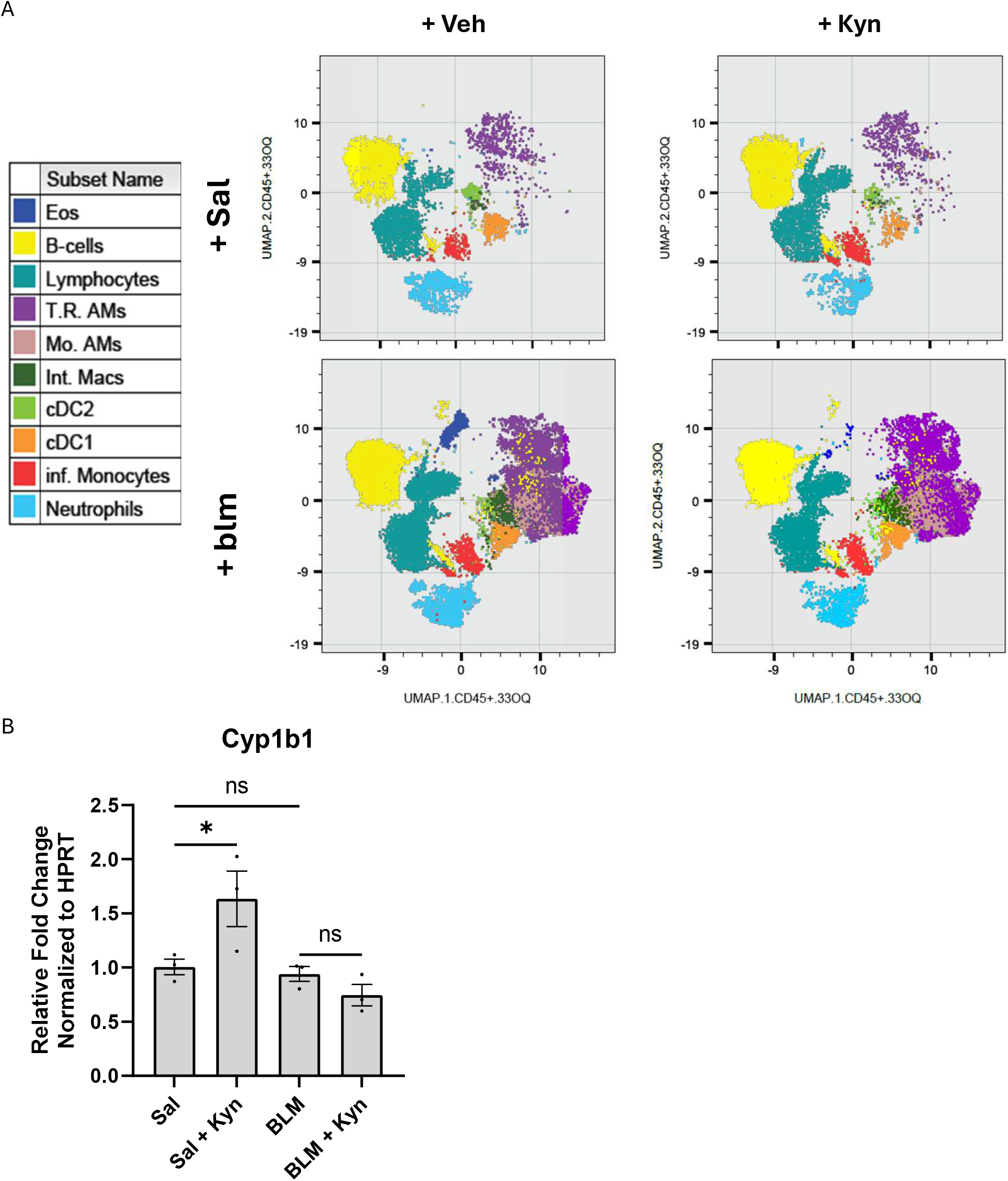
A) Mice were instilled with 1U/kg of bleomycin or saline at day 0, and half were additionally treated with 40mg/kg kynurenine by oral gavage d10-14. Lungs were collected on d14 for collagenase digest and staining. tSNE based on flow cytometry profiles shown here. B) RNA was extracted from whole lung and analyzed for the indicated transcript by qPCR. * = p < 0.05.

**Supplemental Figure 3.**
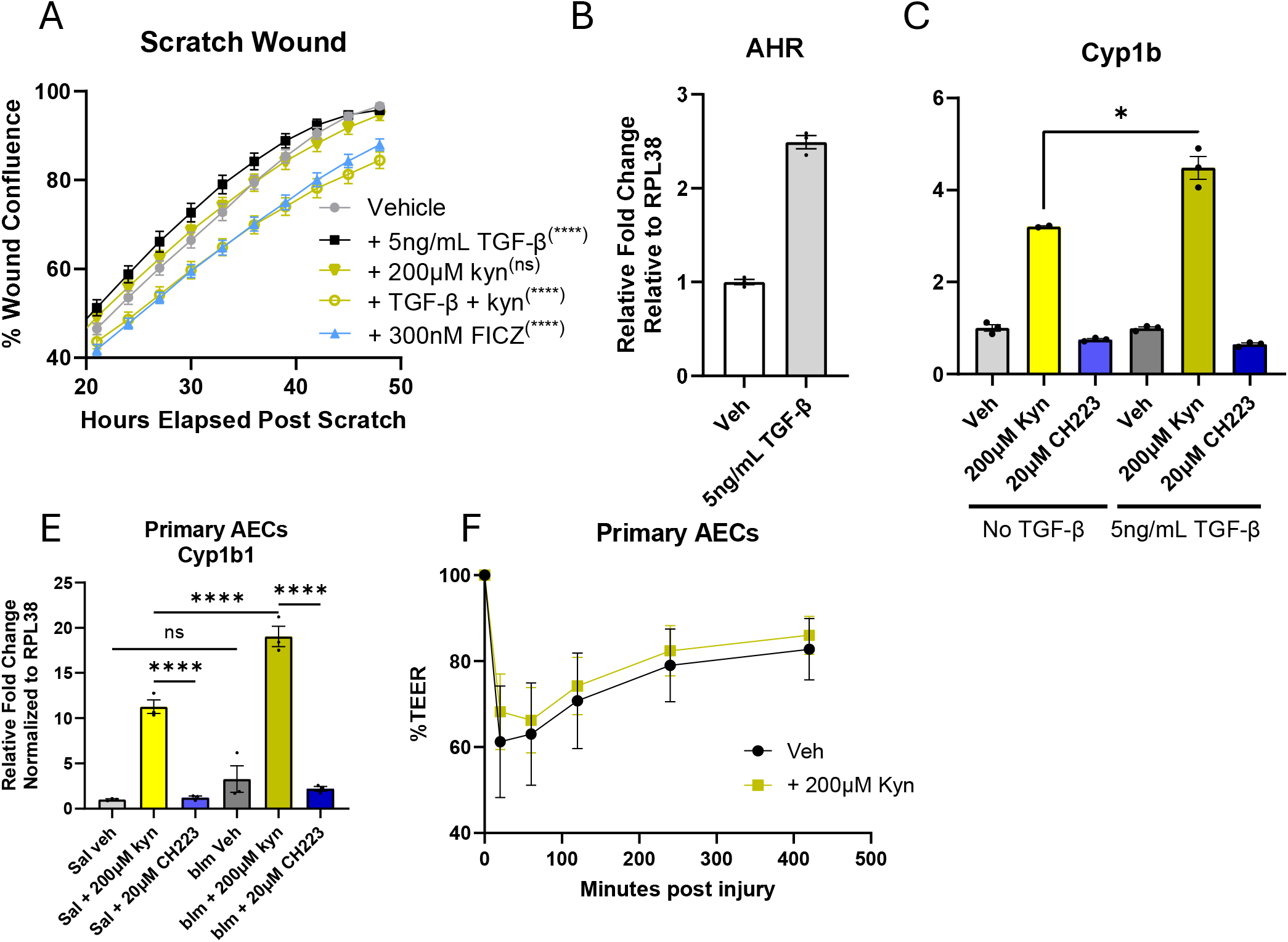
Epithelial cell AHR responses. A) Scratch wound of MLE-12 cells in HITES media with veh or 200µM kyn. B-C) qPCR results of AHR and Cyp1B RNA from MLE-12s plated to 70% confluence and treated with 200uM Kyn, 5ng/mL TGF-β, 20μ,M CH223191 or veh as noted. D) qPCR results of Cyp1B1 RNA from primary AECs treated with veh, kyn, or CH223 overnight. E) Trans-epithelial electrical resistance (TEER) on primary mouse alveolar epithelial cells threated with 100ng/mL TNFα and 100ng/mL IFNγ, and 200uM Kyn as noted in modified HITES media. Error bars SEM. Significant determined by ANOVA. Results representative of at least two experiments. * = p < 0.05, *** = p < 0.001, **** = p < 0.0001, **** = p < 0.0001.

