## Supplementary material for "Cell-Specific Modulation of the Aryl Hydrocarbon Receptor by Kynurenine in Pulmonary Fibrosis Requires Microenvironmental Crosstalk": Table of PCR primers

Table 1

Assay IDs for IDT Smartset qPCR Primers

| **Gene** | **IDT Assay ID Number** |
| --- | --- |
| Mouse IL-6 | Mm.PT.58.10005566 |
| Mouse AHR | Mm.PT.58.12887303 |
| Human AHR | Hs.PT.56a.38998805 |
| Mouse Cyp1b1 | Mm.PT.58.43705524 |
| Human Cyp1b1 | Hs.PT.58.25328727.g |
| Human Col1a1 | Hs.PT.58.15517795 |
| Human ACTA2 (αSMA) | Hs.PT.56a.2542642 |
